# Srrm234, but not canonical SR and hnRNP proteins drive inclusion of *Dscam* exon 9 variable exons

**DOI:** 10.1101/584003

**Authors:** Pinar Ustaoglu, Irmgard U. Haussmann, Hongzhi Liao, Antonio Torres-Mendez, Roland Arnold, Manuel Irimia, Matthias Soller

**Affiliations:** School of Biosciences, College of Life and Environmental Sciences, University of Birmingham, Edgbaston, Birmingham, B15 2TT, United Kingdom; Department of Life Science, School of Health Sciences, Birmingham City University, Birmingham, B5 3TN, United Kingdom; Centre for Genomic Regulation, Barcelona Institute of Science and Technology (BIST), Barcelona 08003, Spain; Institute of Cancer and Genomics Sciences, College of Medical and Dental Sciences, University of Birmingham, Edgbaston, Birmingham, B15 2TT, United Kingdom; Universitat Pompeu Fabra (UPF), Barcelona 08003, Spain; ICREA, Barcelona 08010, Spain

**Keywords:** Alternative splicing, *Dscam*, RNA binding proteins, SR proteins, SRm300, hnRNP proteins

## Abstract

Alternative splicing of pre-mRNA is a major mechanism to diversify protein functionality in metazoans from a limited number of genes. In the *Drosophila melanogaster Down Syndrome Cell Adhesion Molecule (Dscam)* important for neuronal wiring up to 38,016 isoforms can be generated by mutually exclusive alternative splicing in four clusters of variable exons. However, it is not understood how a specific exon is chosen from the many variables and how variable exons are prevented from being spliced together. A main role in the regulation of *Dscam* alternative splicing has been attributed to RNA binding proteins, but how they impact on exon selection is not well understood. Serine-arginine-rich (SR) proteins and hnRNP proteins are the two main types of RNA binding proteins with major roles in exon definition and splice site selection. Here, we analyzed the role of SR and hnRNP proteins in *Dscam* exon 9 alternative splicing in mutant *Drosophila melanogaster* embryos because of their essential function for development. Strikingly, loss or overexpression of canonical SR and hnRNP proteins even when multiple proteins are depleted together, does not affect *Dscam* alternative exon selection very dramatically. Conversely, non-canonical SR protein Serine-arginine repetitive matrix 2/3/4 (Srrm234) is a main determinant of exon inclusion in *Dscam* exon 9 cluster. Since long-range base-pairings are absent in the exon 9 cluster, our data argue for a small complement of regulatory factors as main determinants of exon inclusion in the *Dscam* exon 9 cluster.

## Introduction

During alternative splicing the combination of exons can be varied to generate multiple different transcripts and proteins from one gene (Soller 2006; Nilsen and Graveley 2010; Fiszbein and Kornblihtt 2017). In humans, 95% of genes, and in *Drosophila melanogaster* 63% of genes are alternatively spliced, respectively (Wang et al. 2008; Fu and Ares 2014). Among the genes where alternative splicing generates the greatest diversity of isoforms is the *Drosophila melanogaster* homolog of human *Down Syndrome Cell Adhesion Molecule* (*Dscam*), which encodes a cell surface protein of the immunoglobulin superfamily. The *Dscam* gene comprises 95 alternatively spliced exons that are organized into four clusters, namely 4, 6, 9 and 17 which contain 12, 48, 33 and 2 variables, respectively. Hence, the *Dscam* gene can generate up to 38,016 different proteins (Schmucker et al. 2000; Neves et al. 2004; Hemani and Soller 2012; Sun et al. 2013). *Dscam* is functionally required for neuronal wiring in the nervous system, but also for phagocytosis of invading pathogens in the immune system (Schmucker et al. 2000; Watson et al. 2005). Interestingly, *Dscam* in mosquitos changes its splicing pattern upon pathogen exposure to produce isoforms with higher binding affinity for binding pathogen (Dong et al. 2006). However, despite intense research, relatively little is known about how *Dscam* alternative splicing is regulated in flies.

Pre-mRNA splicing is a multistep process catalyzed by the spliceosome sequentially assembled from five U snRNPs together with numerous proteins. Spliceosome assembly initiates by the recognition of the 5’ splice site by U1 snRNP and of the 3’ splice site by U2 snRNP together with U2AFs recognizing the branchpoint and the polypyrimidine tract and the AG of the 3’ splice site, respectively. Then the U4/5/6 tri-snRNP is recruited and upon several structural rearrangements where U4 snRNP leaves the spliceosome, catalysis takes place by two transesterification reactions (Luhrmann and Stark 2009).

Alternative splicing is to a large degree regulated at the level of splice site recognition involving base-pairing of U1 snRNP to the 5’ splice site YAG/GURAGU and U2 to the branchpoint WNCUAAU (W: A or U, *Drosophila melanogaster* consensus(Lim and Burge 2001)) whereby splice sites closer to the consensus are preferably used (Soller 2006). Splice site selection is critically assisted by RNA binding proteins (RBPs) that support or inhibit recognition of splice sites.

Serine-arginine rich (SR) and heterogenous nuclear ribonucleoproteins (hnRNPs) are two prominent classes of RBPs involved in alternative splicing regulation (Busch and Hertel 2012; Fu and Ares 2014; Bradley et al. 2015). Humans have twelve and flies have eight SR proteins each having one or two RNA Recognition Motifs (RRM) and RS domain rich in serines and arginines (Busch and Hertel 2012). In addition, RS domains are present in some other splicing factors lacking RNA binding domains such as Tra2, SRRM1 (SRm160), and SRRM2 (SRm300), SRRM3, and SRRM4 (nSR100), which are termed non-canonical SR proteins (Blencowe et al. 1999; Long and Caceres 2009; Busch and Hertel 2012; Best et al. 2014). In contrast, hnRNP proteins are more diverse in their modular assembly containing RNA binding domains (e.g. RRM, KH or RGG domains) and various auxiliary domains (Geuens et al. 2016). In humans, the most prominent hnRNPs are the abundantly expressed hnRNP A and C family (Busch and Hertel 2012; Geuens et al. 2016).

SR proteins mostly bind to exonic splicing enhancers (ESEs) and recruit spliceosomal components to splice sites through their RS domains. SR protein binding sites are present in both alternatively spliced and constitutive exons to promote exon inclusion, but they can also repress inclusion of alternative exons (Black 2003; Shen et al. 2004; Wang et al. 2006; Chen and Manley 2009; Pandit et al. 2013). Although SR proteins recognize similar sequences, distinct functions have been shown either by binding distinct sites or when bound to the same site through differential regulation mediated by combinatorial interactions with other splicing regulators (Gabut et al. 2007; Anko et al. 2012; Pandit et al. 2013; Bradley et al. 2015).

In contrast to SR proteins, hnRNP proteins mostly bind to intronic splicing silencers (ISSs) and repress inclusion of alternative exons (House and Lynch 2006; Wang et al. 2008). In addition, they can act antagonistically to SR proteins by binding to exonic splicing silencers (ESSs) and compete with SR proteins for binding (Soller 2006; Long and Caceres 2009). However, a more comprehensive analysis revealed that SR and hnRNP proteins can also act coordinately in many instances in exon inclusion or repression (Brooks et al. 2015).

With regard to the alternative splicing in the *Dscam* gene a model has been proposed for the exon 6 cluster involving long-range base-pairing. Here, a conserved docking sequence in the first intron of the exon 6 cluster can base-pair with complementary selector sequences in front of each variable exon to bring a chosen variable exon into the proximity of the proximal constant flanking exon for splicing (Graveley 2005). This model also requires that the entire cluster is maintained in a repressed state and variable exons are selected under the control of RBPs. However, the architecture in the exon 4 an 9 cluster is different and conserved “docking site” sequences are found at the end of the exon 4 and 9 clusters, but the support for this model based on evolutionary sequence conservation is weak (Yang et al. 2011; Haussmann et al. 2019).

RNAi knock-down of RBPs in cell culture in *Drosophila melanogaster* S2 cells revealed little changes for most variable exons in the *Dscam* exon 4 cluster, but this result could be due to residual protein left (Park et al. 2004). Hence, we wanted to investigate the role of SR and hnRNP proteins in *Drosophila melanogaster Dscam* alternative splicing more comprehensively at an organismal level using knock-out mutants and overexpression. Most SR and hnRNP proteins are essential for development to adult flies, but embryonic development proceeds such that *Dscam* alternative splicing could be analysed in late stage embryos in SR and hnRNP mutants or when over-expressed. Unexpectedly, we find that inclusion of *Dscam* exon 9 variables is affected little in loss or gain of function conditions. Likewise, even upon removal even of multiple factors *Dscam* exon 9 alternative splicing changes little. However, the non-canonical SR protein Serine-arginine repetitive matrix 2/3/4 (Srrm234) is required for inclusion of most exon 9 variables. We further find that long-range base-pairing is not supported as a general model. Hence, our results argue that a small complement of RBPs are main regulators of *Dscam* exon 9 alternative splicing.

## Results

### Analysis of *Dscam* exon 9 alternative splicing by restriction digests

All 33 exons in the *Drosophila melanogaster Dscam* exon 9 variable exon cluster have about the same length and run as a single band on an agarose gel (Figure 1A and B). The sequences in variable exons, however, differ enough such that a complement of restriction enzymes can digest the complex mix of PCR products to identify a majority of isoforms on sequencing type gels using one ^32^P labelled primer after reverse transcription of the mRNA (Figure 1C and D) (Haussmann et al. 2019). Using a combination of SacII, ClaI, PshAI, HaeIII, MseI, BsrI and BstNI yields 26 fragments of unique size identifying 20 variable exons (Fig 1D).

**Figure 1.**
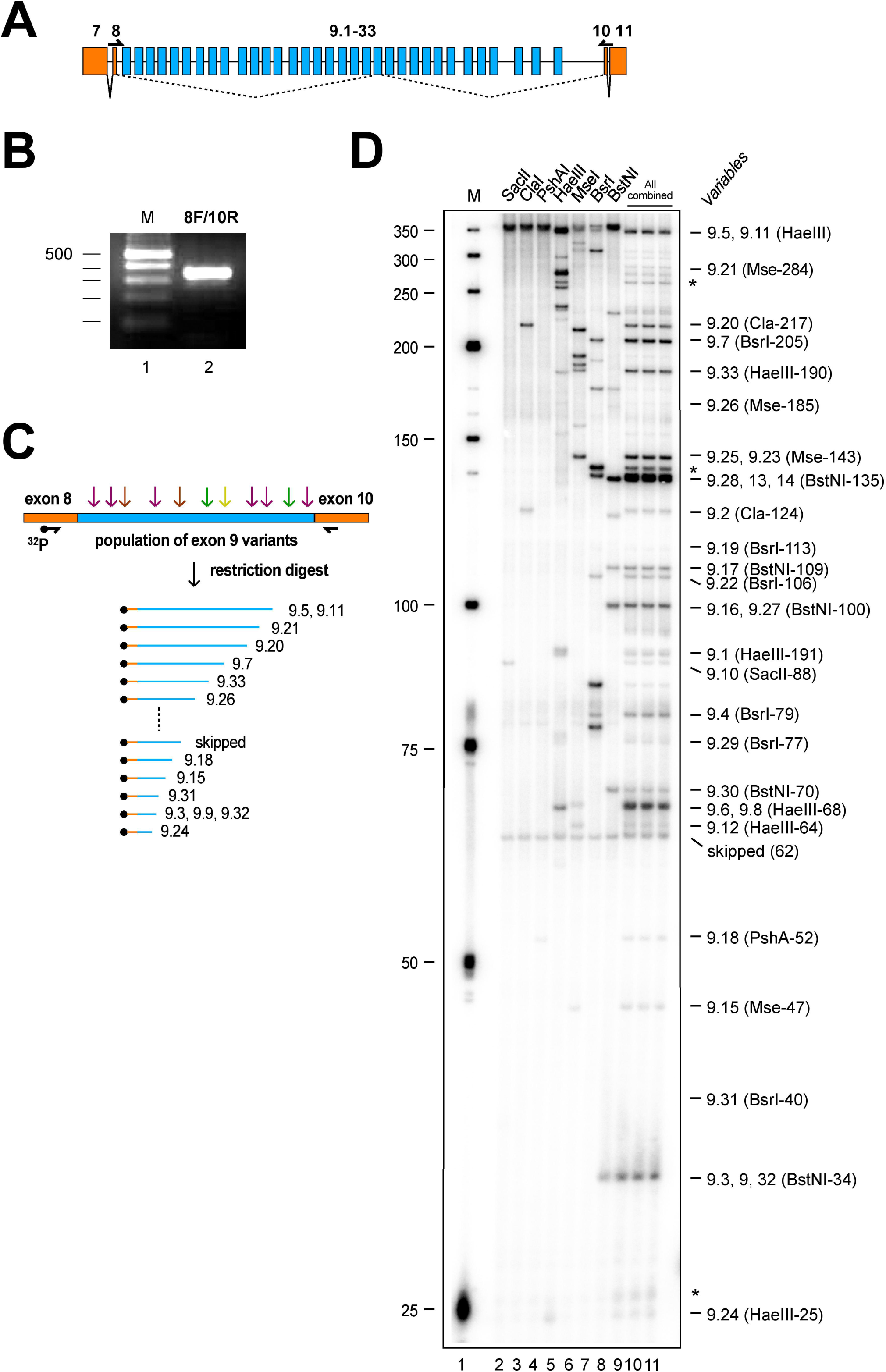
Analysis of *Dscam* exon 9 alternative splicing by restriction digestion of PCR products. (A) Schematic of the *Dscam* exon 9 variable cluster gene region. Constitutive exons are shown in orange and variable exons in blue. Primers to amplify the variable part are shown on top of the exons. (B) RT-PCR product for variable exon cluster 9 shown on a 3% agarose gel. (C) Schematic of the method used to resolve inclusion levels of variable exons using a ^32^P labelled forward primer in combination with a set of restriction enzymes followed by separation of a denaturing polyacrylamide gel. (D)Denaturing acrylamide gel (6%) showing a restriction digest (SacII, ClaI, PshAI, HaeIII, MseI, BsrI, BstNI) of *Dscam* exon 9 variables amplified with a ^32^P labelled forward primer from 14-18 h *Drosophila* embryos. Single enzyme reference digests are shown on the left (lanes 2-8) and the combination of all enzymes on the right (lanes 9-11).

### Alteration of SR proteins has little impact on *Dscam* exon 9 alternative splicing

SR proteins are organized into five families and representative orthologues are present in *Drosophila melanogaster* (Fig 2) (Busch and Hertel 2012). To determine the role of SR proteins in *Drosophila melanogaster Dscam* exon 9 alternative splicing we obtained mutants and over-expression lines for most of the canonical SR proteins as well as for general splicing factor SF1 and non-canonical SR protein Srrm1 (SRm160) and Srrm234 (SRm300, CG7971) (Fig 2A, Supplementary Fig S1). For loss of function (LOF) alleles of SR genes, five from the ten analysed, which are *X16^GS1678^, SF2^GS22325^, B52^29^, Srrm1 ^B103^* and *Srrm234*^Δ*N*^, are required to reach adulthood (Fig 2A and B). Gain of function (GOF) conditions by pan-neural *elavGAL4* mediated over-expression via *UAS* was lethal in larval instars for *UAS GFP-X16, UAS RSF1, UAS GFP-SC35, UAS GFP-SF2* and *UAS GFP-B52*, while over-expression of *UAS SF1* and *UAS Srrm1* from *EP* lines did not result in a phenotype (Fig 2A and C).

**Figure 2.**
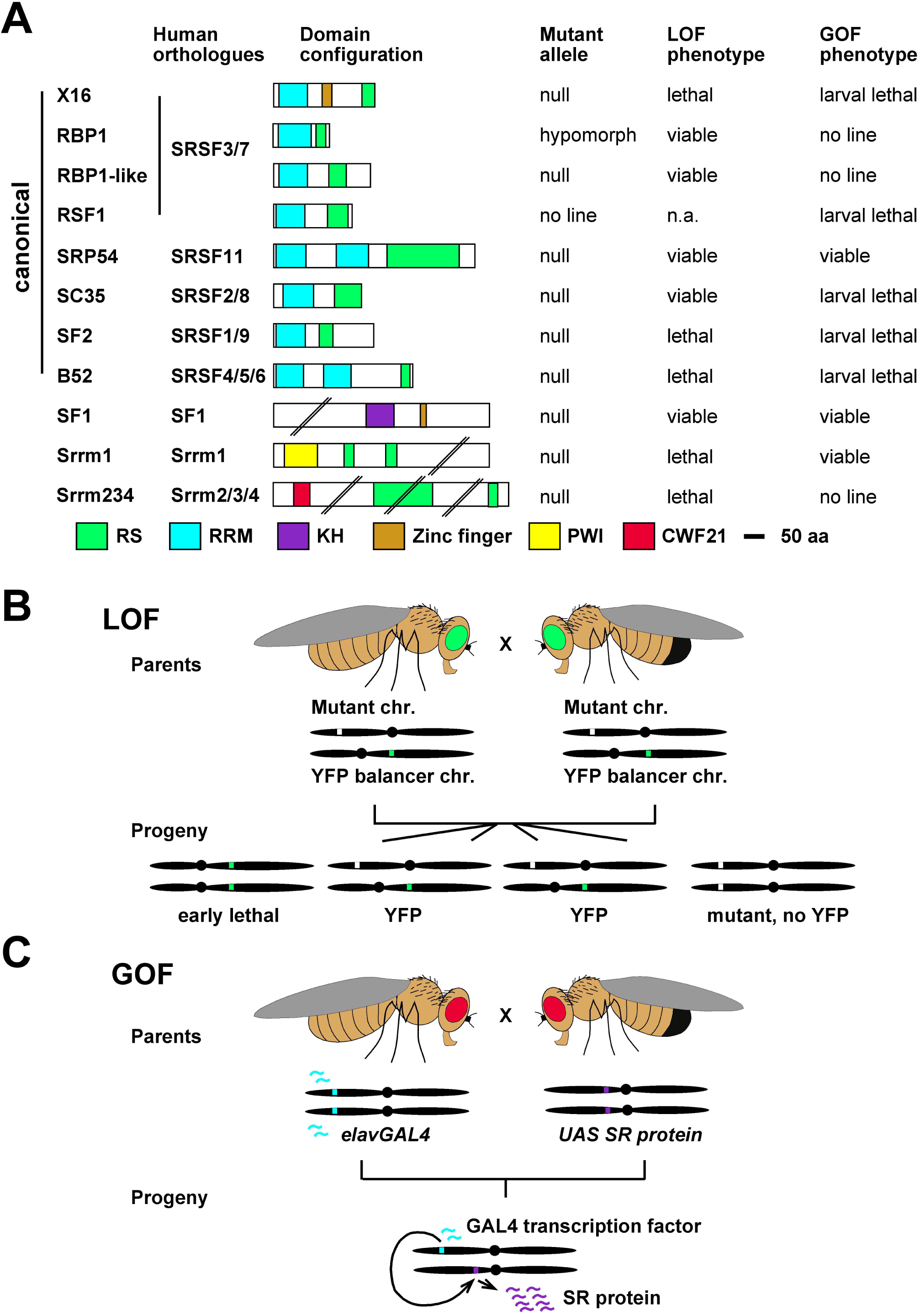
Protein domain structure of *Drosophila* SR proteins and phenotype of loss of function (LOF) and gain of function (GOF) mutants. (A) Evolutionary relationship of *Drosophila* SR proteins with human homologues is shown on the left and the domain structure is indicated by coloured boxes. Arginine-serine-rich domain (RS, green), RNA recognition domain (RRM, light blue), hnRNP K homology domain (KH, purple), zinc finger domain (ochre), Proline-Tryptophan-Isoleucine domain (PWI, yellow) and CWF21 domain (red). The type of allele obtained, and the loss (LOF) and gain of function (GOF) phenotypes are indicated on the right. (B) Viability was determined from stocks that harbour a zygotically expressing GFP marked balancer chromosome, which contains a set of inversions to supress recombination and a recessive lethal mutation. If a gene is essential, only heterozygous flies will survive. Homozygous mutant embryos were identified in the progeny of these stocks, by the lack of GFP and advanced development as homozygous balancer carrying embryos die early before GFP expression. (C) To obtain embryos overexpressing SR proteins flies carrying the yeast GAL4 transcription factor under the control of the pan-neuronal *elav* promoter were crossed with lines harbouring SR proteins under the control of the yeast *UAS* promoter.

Next, we analysed inclusion levels of exon 9 variables in embryos for LOF and GOF conditions of canonical SR proteins and SF1. For LOF alleles *X16 ^GS1678^, RBP1 ^HP37044^, RBP1-like ^NP0295^, Srp54^GS15334^, SC35^KG02986^, SF2^GS22325^*, *B52^28^* and *SF1^G14313^* the *Dscam* exon 9 splicing pattern unexpectedly remained largely unchanged and significant changes are prominent in exon 9.4, 9.21 and 9.5/9.11 for mutants compared to wild type (marked in red for decrease and blue for increase, Figure 3A and B). Similar results were obtained for over-expression of *UAS GFP-X16, UAS RSF1, UAS GFP-SC35, UAS GFP-SF2, UAS GFP-B52* and *UAS-Srrm1* with significant changes only prominent in exon 9.19 and 9.21 for GOF conditions compared to wild type (marked in red for decrease and blue for increase, Fig 3C and D).

**Figure 3.**
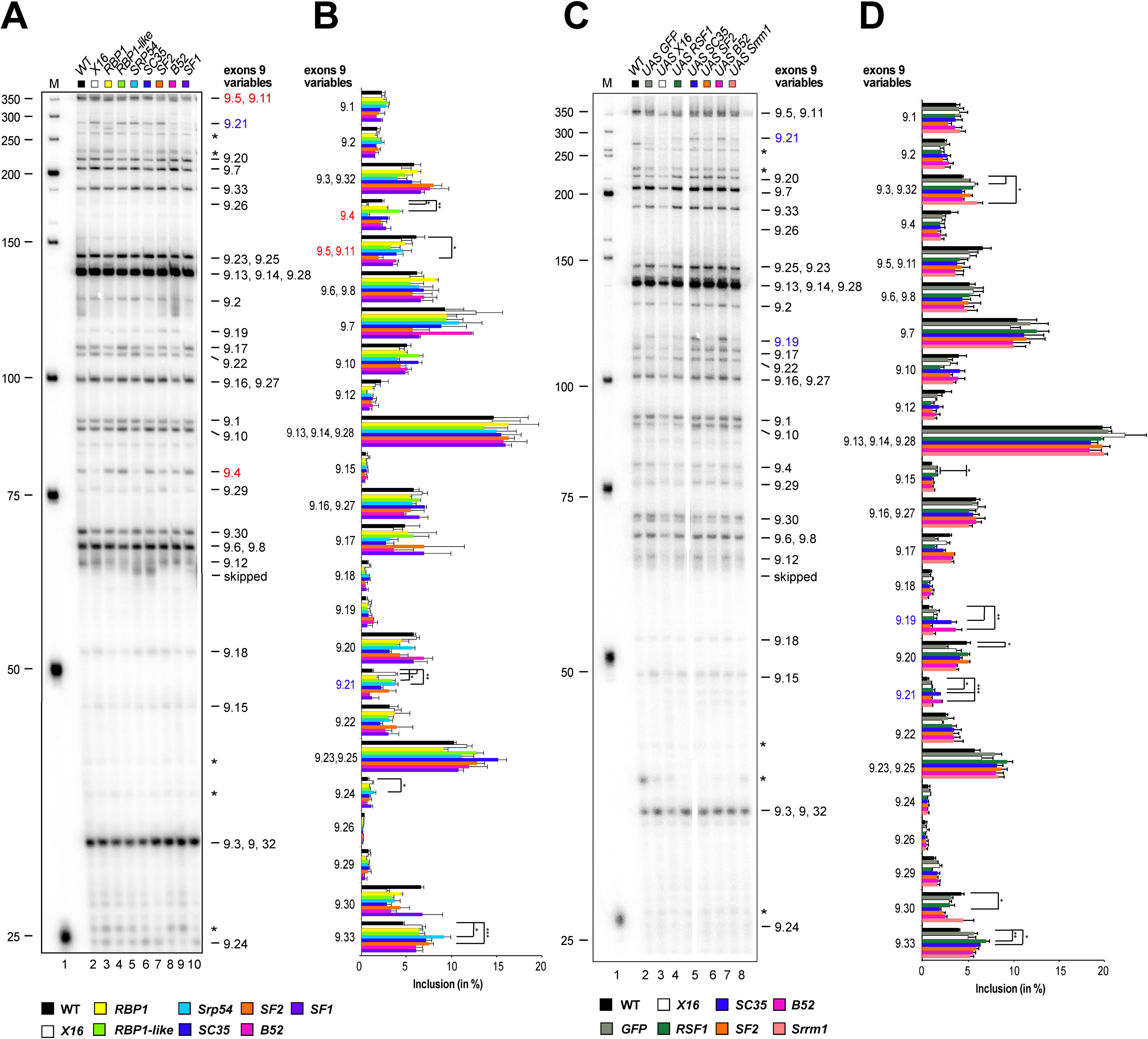
Analysis of *Dscam* exon 9 alternative splicing in canonical SR protein loss of function (LOF) and gain of function (GOF) mutants. (A, C) Denaturing acrylamide gel showing restriction digests of *Dscam* exon 9 variables amplified with a ^32^P labelled forward primer from 14-18 h *Drosophila* embryos for canonical SR protein LOF (A) and *elavGAL4 UAS* GOF mutants (C). Quantification of inclusion levels are shown as means with standard error from three experiments for canonical SR protein LOF (B) and GOF mutants (D). Prominent changes in inclusion levels in mutants compared to wild type are marked with red letters for a decrease and in blue for an increase, and statistically significant differences are indicated by asterisks (***p ≤ 0.001, **p ≤ 0.01, *p ≤ 0.05).

### Non-canonical SR protein Srrm234 is required for selection of *Dscam* exon 9 variables

The canonical SR proteins contain an RRM and bind to RNA. In contrast, the large Srrm1 and Srrm234 proteins contain RS domains, but seem not to bind RNA and exert splicing enhancing functions through association with proteins bound to ESEs (Fig 2) (Blencowe et al. 1998; Eldridge et al. 1999; Blencowe et al. 2000; Szymczyna et al. 2003).

We obtained null mutants for both genes, *Srrm1 ^B103^* and *Srrm234*^Δ*N*^, which are late embryonic lethal (Fan et al. 2014). While loss of Srrm1 had little effect on *Dscam* exon splicing, loss of Srrm234 resulted in significant reduction in inclusion for many variable exons (marked in red, 9.4, 9.7, 9.10, 9.16/9.27, 9.17, 9.18, 9.20, 9.23/9.25, 9.29 and 9.33) that is compensated by increased inclusion of a few variable exons (marked in blue, 9.3/9.32, 9.6/9.8, 9.12 and 9.30) (Figure 4A and B). Consistent with a role in *Dscam* exon 9 alternative splicing regulation, *Srrm234* is also expressed in the nervous system of *Drosophila melanogaster* embryos in the same pattern as *Dscam* (Supplementary Fig S2).

**Figure 4.**
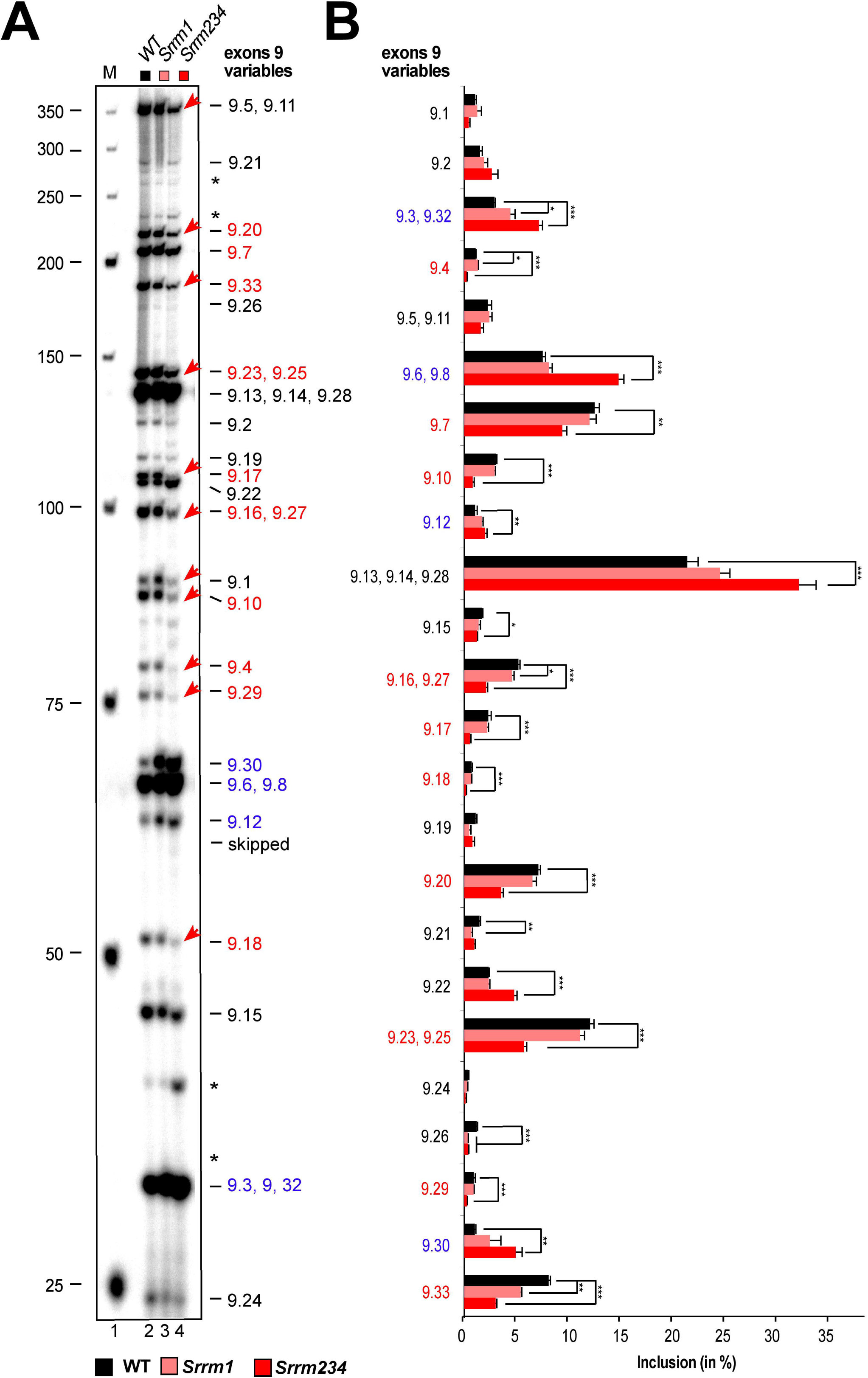
Analysis of *Dscam* exon 9 alternative splicing in non-canonical SR proteins Srrm1 and Srrm234. (A) Denaturing acrylamide gel showing restriction digests of *Dscam* exon 9 variables amplified with a ^32^P labelled forward primer from 14-18 h *Drosophila* embryos for Srrm1 and Srrm234 protein LOF mutants. Quantification of inclusion levels are shown as means with standard error from three experiments (B). Red arrows point towards exons with reduced inclusion levels in the *Srrm234*^Δ^*^N^* mutant compared to wild type. Prominent changes in inclusion levels are marked with red letters for a decrease and in blue for an increase in the *Srrm234*^Δ^*^N^* mutant compared to wild type. Statistically significant differences are indicated by asterisks (***p ≤ 0.001, **p ≤ 0.01, *p ≤ 0.05).

### Alteration of hnRNP proteins has little impact on *Dscam* exon 9 alternative splicing

Since alterations of individual canonical SR proteins had little impact on selection of *Dscam* exon 9 variables, we focused on hnRNP proteins as candidates for repressing inclusion of exon 9 variables (Olson et al. 2007; Chen and Manley 2009; Fu and Ares 2014). *Drosophila melanogaster* has four members of the highly expressed hnRNP A family (Hrp36, Hrp38, Rb97D and Hrp48) and one member of the hnRNP C, although the *Drosophila melanogaster* orthologue is considerably longer (Fig 5)(Appocher et al. 2017). Other highly expressed hnRNP proteins are Hrp40 (hnRNP D), Glorund (hnRNP F/H), Hrb57A (hnRNP K) and Hrb59 (hnRNP M).

**Figure 5.**
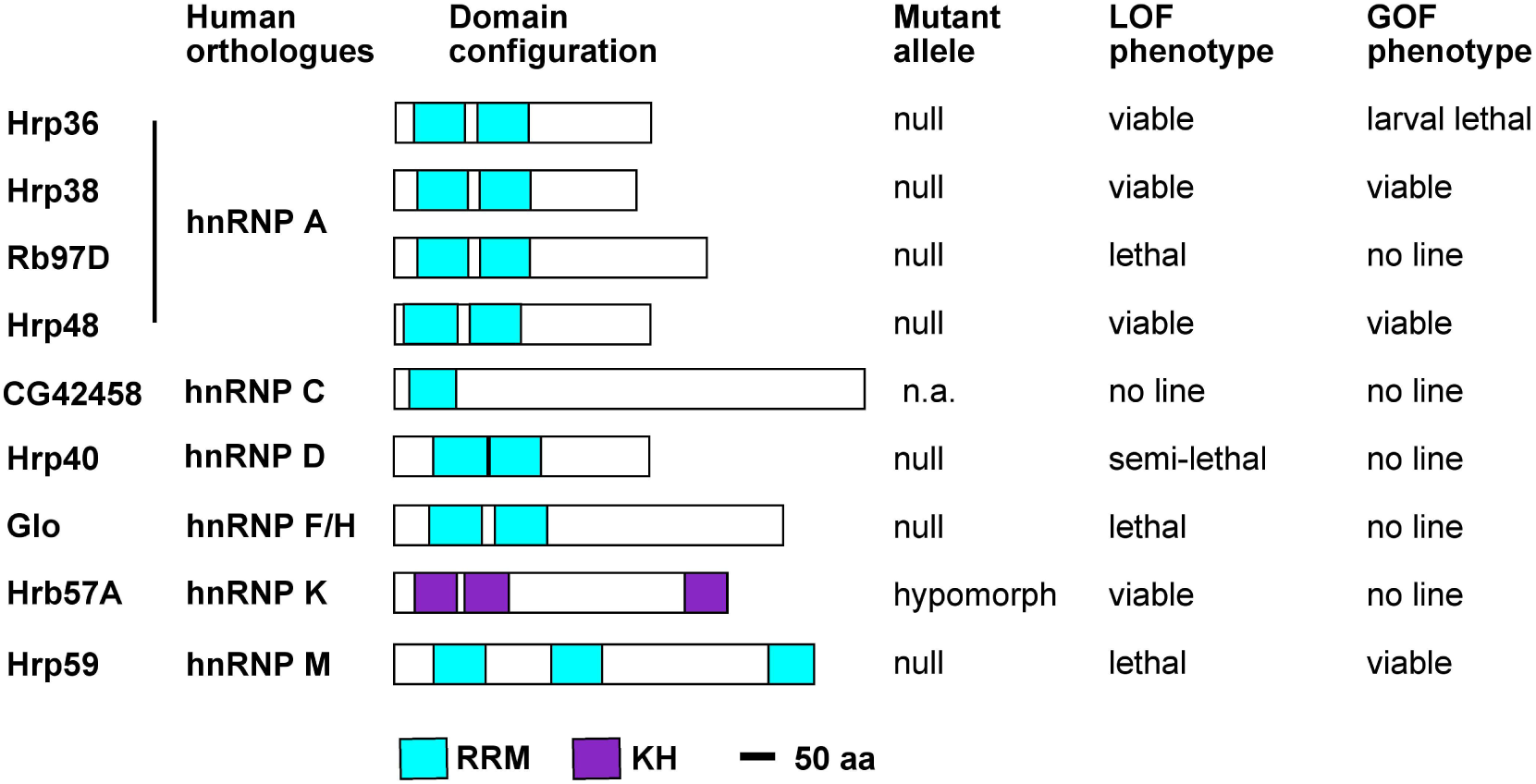
Protein domain structure of *Drosophila* hnRNP proteins and phenotype of loss of function (LOF) and gain of function (GOF) mutants. Evolutionary relationship of *Drosophila* hnRNP proteins with human homologues is shown on the left and the domain structure is indicated by coloured boxes. RNA recognition domain (RRM, light blue) and hnRNP K homology domain (KH, purple). The type of allele obtained, and the loss (LOF) and gain of function (GOF) phenotypes are indicated on the right.

To determine the role of hnRNP proteins in *Drosophila melanogaster Dscam* exon 9 alternative splicing we could obtain null-mutants for all major hnRNP proteins (*Hrp36^BG02743^, Hrp38^MI1059^, RB97D^1^*, *Hrp48 ^GS14498^, Hrp40 ^GS18188^, glo ^f02674^, Hrb57 ^G13574^* and *Hrp59^GS6029^*) and gene-switch or *EP* over-expression lines for most (Hrp36 *^GS15926^*, Hrp38 *^GS12795^*, Hrp48 *^EY12571^*, and Hrp59 *^GS6029^*, Fig 5, Fig 7A, Supplementary Fig S3). Half of the tested major hnRNPs are required for viability (*Rb97D*, *Hrp40*, *glo* and *Hrp59*), while only over-expression of Hrp36 was lethal (Fig 5).

Then, we analysed *Dscam* exon 9 inclusion levels for LOF and GOF of hnRNPs. For LOF alleles, the *Dscam* exon 9 splicing pattern also unexpectedly remained largely unchanged and significant changes are prominent in exon 9.17, 9.19 and 9.21 for mutants compared to wild type (marked in red for decrease and blue for increase, Fig 6A and B). Similar results were obtained for GOF conditions with significant changes only prominent in exon 9.2, 9.4, 9.17, 9.19, 9.21, and 9.30 (marked in red for decrease and blue for increase, Fig 6C and D).

**Figure 6.**
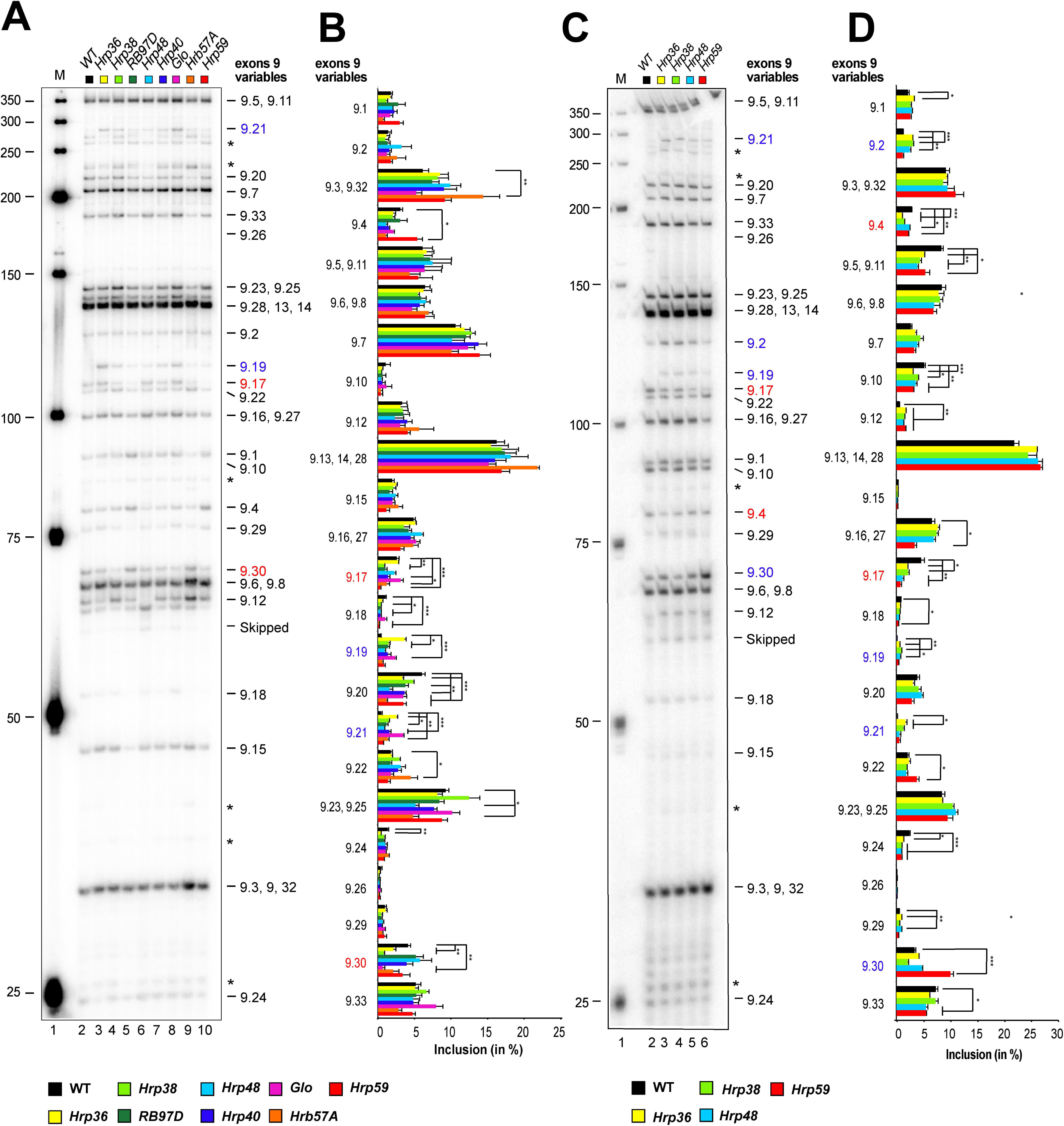
Analysis of *Dscam* exon 9 alternative splicing in general hnRNP protein loss of function (LOF) and gain of function (GOF) mutants. (A, C) Denaturing acrylamide gel showing restriction digests of *Dscam* exon 9 variables amplified with a ^32^P labelled forward primer from 14-18 h *Drosophila* embryos for general hnRNP protein LOF (A) and *elavGAL4 UAS* GOF mutants (C). Quantification of inclusion levels are shown as means with standard error from three experiments for canonical SR protein LOF (B) and GOF mutants (D). Prominent changes in inclusion levels are marked with red letters for a decrease and in blue for an increase, and statistically significant differences are indicated by asterisks (***p ≤ 0.001, **p ≤ 0.01, *p ≤ 0.05).

Binding of Hrp36, Hrp38, Hrp48 and Hrp40 to RNA has been analysed globally in *Drosophila melanogaster* and binding motifs have been established by SELEX (Blanchette et al. 2009). When we reanalysed these data we did find significantly increased binding for all four Hrp proteins (p<0.05) to the *Dscam* exon 9 variable cluster compared to averaged binding, but the binding curves did not overlap with changes in exon inclusion in LOF or GOF conditions (Supplementary Figure S4). Likewise, we did not find an overlap of SELEX motif enrichments with changes in exon inclusion in LOF or GOF conditions (Supplementary Figure S4).

### Removal of multiple SR and hnRNP proteins has little impact on selection of *Dscam* exon 9 variables

hnRNP36 and SR protein B52 genes lie next to each other in the *Drosophila melanogaster a* genome and potentially could cross-regulate to compensate for each other (Fig 7A). Therefore, we generated a double knock-out of *hnRNP36* and *B52* genes (*hnRNP36/B52*^Δ^*^1^*). In addition, we also combined this double knock-out with the *hnRNP38* mutant as *hnRNP36* and *hnRNP38* are closely related and since they are highly expressed, they could act redundantly.

**Figure 7.**
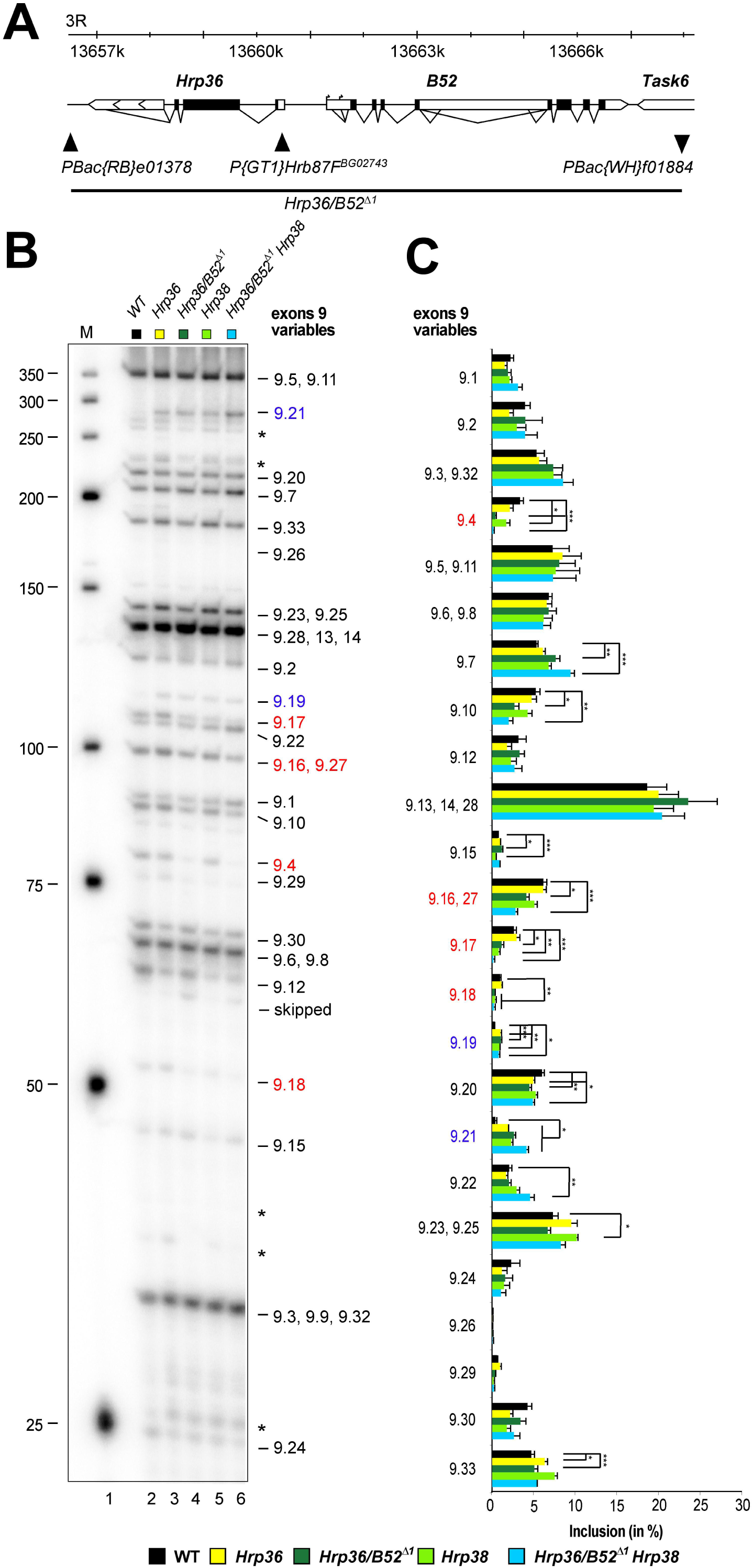
Analysis of *Dscam* exon 9 alternative splicing in double and triple mutants for general splicing regulators. (A) Schematic of the genomic region of *Hrp36* and *B52* genes. Transposon inserts are shown with triangles and the deletion Hrp36/B52^Δ^*^1^* is indicated at the bottom. (B) Denaturing acrylamide gel showing restriction digests of *Dscam* exon 9 variables amplified with a ^32^P labelled forward primer from 14-18 h *Drosophila* embryos for Hrp36^BG02743^, Hrp36/B52^Δ*1*^, Hrp38^d05172^ and Hrp36/B52^Δ*1*^ Hrp38^d05172^ mutants. (C) Quantification of inclusion levels are shown as means with standard error from three experiments. Prominent changes in inclusion levels are marked with red letters for a decrease and in blue for an increase, statistically significant differences are indicated by asterisks (***p≤ 0.001, **p≤ 0.01, *p≤ 0.05).

Surprisingly, even in *hnRNP36/B52*^Δ1^*hnRNP38*^*d05172*^ triple mutants, *Dscam* exon 9 was robustly spliced with only differences in inclusion levels of variables 9.4, 9.16/9.27, 9.17, 9.18, 9.19, and 9.21 compared to controls (marked in red for decrease and blue for increase, Fig 7B and C.

### Variable exon selection is not explained by long-range base-pairing in *Dscam* exon 9

In the exon 4 cluster, we could not detect conserved sequences adjacent to every variable exon that could mediate long-range base-pairing arguing that such a mechanism is not involved in variable exon 4 selection (Haussmann et al. 2019). A sequence alignment between *Drosophila melanogaster* and *Drosophila virilis* showed strong conservation in the coding sequences and the architecture of the exon 9 cluster with only few insertions and deletions of exons (Supplementary Fig S5). There are conserved sequence elements in the intron before constant exon 10 that potentially could serve as a docking site (Yang et al. 2011), but we did not find conserved sequence elements between every variable exon by manual inspection (Supplementary Fig S6). A systematic bioinformatics comparison of this potential docking site with arbitrary sampled sequences of the same length and sequence complexity further did not reveal a special propensity of the docking site to form more or stronger complementary alignments within *Dscam* intronic sequences in the variable exon 9 cluster. The same picture arises when comparing predicted energies of the best secondary duplex structures formed by the reverse intronic sequence, the docking site, and sampled sequences (Supplementary Fig S7). In addition, we didn’t find other instances of the docking site in any other gene of *D. melanogaster* (except in the anti-sense RNA *CR45129* of *Dscam*), indicating that the sequence is not recurrent in other genes.

## Discussion

Although the sequence determinants that direct the spliceosome to its correct position are very degenerate, splicing needs to occur precisely and with high accuracy to prevent disease (Cooper et al. 2009; Zaharieva et al. 2012). It is therefore thought that RNA binding proteins play key roles in localizing functional splice sites. In particular, the abundant SR and hnRNP proteins have been attributed key roles in this process by forming RNA-protein complexes co-transcriptionally to recruit early splicosomal components for defining splice sites. Although SR proteins were initially viewed as binding ESEs to activate splicing, and hnRNP proteins to bind ISSs for antagonizing SR proteins, a number of genome-wide studies draw a more complex picture for both SR and hnRNP binding and function (Blanchette et al. 2009; Anko et al. 2010; Anko et al. 2012; Huelga et al. 2012; Pandit et al. 2013; Brooks et al. 2015). In fact, both SR and hnRNP proteins can have very specific functions in one context, but also redundant functions in another context.

In this regard, we hypothesised that an array of similar exons as found in the *Drosophila melanogaster Dscam* gene would provide a platform for SR and hnRNP proteins to evolve exon specific functions to regulate their inclusion. Surprisingly, however, *Dscam* exon 9 alternative splicing is exactly the opposite and the splicing pattern is very robustly maintained when SR and hnRNP proteins were either removed or overexpressed. In particular, the advances of *Drosophila melanogaster* genetics allowed us to use complete knock-outs of most canonical SR and general hnRNP proteins and thus avoid the ambiguity of RNAi that would leave residual protein. Hence, despite complete loss of individual SR and hnRNP proteins, or combinations thereof, the *Dscam* splicing pattern is robustly maintained.

Two explanations are possible for this scenario. First, SR and hnRNP proteins act redundantly at the very extreme such that fluctuations in many would need to occur to impact on *Dscam* alternative splicing. However, whether this model applies will be difficult to test as removal of many general splicing factors will likely lead to global perturbations affecting many genes.

A second scenario could be that *Dscam* alternative splicing is fairly independent of general splicing factors. This would imply a more specific mechanism. Initially, it has been though that long-range base-pairing would provide such a mechanism as conserved sequences have been found in the *Dscam* exon 6 cluster (Graveley 2005). Our previous analysis of the exon 4 cluster, and the in depth analysis of the exon 9 cluster in this paper, however, rule out such mechanism in these two clusters (Haussmann et al. 2019). Hence, the question remains whether two independent mechanisms arose to regulate mutually exclusive alternative splicing in *Dscam* variable clusters. Potentially, the conserved sequences present in the variable clusters could provide binding sites for RBPs that have adopted cluster specific roles. In this context it is interesting to note that deletion of the docking site in exon 6 leads to inclusion of mostly the first exon in the cluster (May et al. 2011). This is unexpected as removal of the splicing activating mechanism should result in skipping of the entire variable cluster, because the proposed repressor hnRNP36 would still be present. Accordingly, the docking site in the exon 6 cluster also exerts a repressive role in maintaining the entire cluster in a repressed state.

Likewise, our finding that non-canonical Srrm234 regulates inclusion of many variables in the *Dscam* exon 9 cluster suggests a mechanism in *Dscam* mutually exclusive alternative splicing that differs from more general splicing rules directed by canonical SR and general hnRNP proteins. One of the human homologs of Srrm234, SRRM4 has key roles in the regulation of microexons (Irimia et al. 2014). Due to their small size, microexons cannot be defined through standard mechanism of exon definition. Hence, a distinct mechanism must apply, that can accurately direct splicing of microexons. Because microexons are often found in large introns, a robust process must underlie their selection and involves a newly described enhancer of microexons domain (eMIC), present in human SRRM3 and SRRM4. Intriguingly, in vertebrates, the ancestral pro-homologue Srrm234 has duplicated into three genes to adopt distinct functions through dedicated protein domains, but these features are maintained by alternative mRNA processing in *Drosophila melanogaster* Srrm234 to include the eMIC at the C-terminus of the protein in neuronal tissue (Torres-Mendez et al. 2019).

*Dscam* alternative exons comply with the general average length of exons and thus the mechanism of their regulation is likely distinct from microexons. *Dscam* exon 9 cluster regulation by Srrm234 seems to involve its Cwf21 domain, which is not required for microexon inclusion (Torres-Mendez et al. 2019). Transposon inserts in the middle of the *Srrm234* gene resulting in a truncated protein, that contains the Cwf21 domain do not affect exon 9 diversity (*Mi{ET1}CG7971^MB07314^* and *Mi(Liang et al.)CG7971^MI04068^*, data not shown). The Cwf21 domain, which is homologous to the yeast Cwc21 domain, has been attributed key roles in splicing due to co-purification of the human ortholog SRRM2 with active spliceosomes and its localization in the catalytic centre of the spliceosome (Bessonov et al. 2008). Furthermore, the Cwc21 domain interacts with the U5 snRNP core components Snu114 and Prp8 involved in key structural rearrangements in the spliceosome during catalysis (Grainger et al. 2009; Gautam et al. 2015). Interestingly, Cwc21 has been attributed roles in splicing of meiotic genes which are regulated differently of general intron containing genes (Gautam et al. 2015). Srrm2 forms a complex with Srrm1 to promote alternative splicing of *Drosophila melanogaster doublesex* required for sex determination, but the mechanism in *Dscam* is different as loss of Srrm1 does not impact on alternative splicing in the exon 9 cluster (Blencowe et al. 1998; Eldridge et al. 1999). In addition, Srrm234 also interacts with U1 70K, so potentially could act to activate 5’ splice site and initiate the splicing process from a repressed state of the variable cluster (Guruharsha et al. 2011).

A *Dscam* cluster-specific role has been suggested for Hrp36 acting as a repressor preventing splicing together of exon 6 variables in S2 cells, but whether the results are the same in flies remains to be tested (Olson et al. 2007). Such a factor has not been identified for exon 4 and 9 clusters, and for the factors tested, we did not observe splicing together of variable exons. Intriguingly, a newly annotated exon 4.0 in the beginning of the bee *Dscam* exon 4 cluster is spliced to exon 4.6 (Decio et al., 2019). This unexpected finding, however, is not compatible with the model described for Hrp36 in the exon 6 cluster, but might involve a repressive sequence element around exon 4.0 similar the element discovered in the beginning of the exon 6 cluster, which when deleted results only in inclusion of exon 6.1 (May et al. 2011). Likewise, factors like Srrm234 would then act as activators to drive inclusion of variables.

Taken together, *Dscam* exon 9 mutually exclusive alternative splicing is robust against fluctuations of in canonical SR and general hnRNP proteins arguing for a specific mechanism regulating inclusion levels of variable exons. Indeed, non-canonical SR protein Srrm234 plays a key role increasing inclusion of many exon 9 variables. However, since Srrm234 does not have not have one of the classic RNA binding domains, additional RNA binding proteins likely connect Srrm234 to *Dscam* exon 9. Hence, our data obtained from knock-outs of general splicing factors indicate that a small complement of RNA binding proteins are likely key regulators of *Dscam* mutually exclusive alternative splicing.

The gene structure of invertebrate *Dscam* harbouring arrays of variable exons for mutually exclusive splicing is at the extreme, but arrays of several alternative exons are common to many genes in metazoans. Likely, a yet to be discovered feature of the spliceosome has been exploited in mutually exclusive alternative splicing of *Dscam* such that only on exon is chosen. Likewise, since sequences that look like splice site are common in large introns, such a mechanism could be broadly relevant for robust select of isolated exons.

## Materials and Methods

### Fly genetics

Flies were maintained on standard cornmeal agar food as described (Haussmann et al. 2013). *CantonS* was used as a wild type control. The following loss of function mutants were used as depicted in Supplementary Figs S1 and S2, and Fig 7A: *X16* (*P{GSV6}Hrb27C^GS16784^,* DGRC (Kyoto #206763*), RBP1 (P{EPg}mRpL37^HP37044^,* BDSC #22011*), RBP1-like (P{GawB}Rbp1-like^NP0295^,* DGRC Kyoto #103580*), Srp54 (P{GSV6}Srp54^GS15334^*, DGRC Kyoto #206174), *SC35 (P{SUPor-P}SC35^KG02986^,* BDSC #12904*), SF2 (P{GSV7}SF2^GS22325^,* DGRC Kyoto #203903*), B52^28^(Gabut et al. 2007), SF1 (P{EP}SF1^G14313^* BDSC #30203*), Hrp36 (P{GT1}Hrb87F^BG02743^,* BDSC #12869*), Hrp38 (Mi(Liang et al.)Hrb98DE^MI10594^,* BDSC #55509*), Rb97D (P{PZ}Rb97D^1^,* BDSC #11782*), Hrp48 (P{GSV6}Hrb27C^GS14498^,* DGRC Kyoto #205836*), Hrp40 (P{GSV6}sqd^GS18188^,* DGRC Kyoto #201020*), Glo (PBac{WH}glo^f02674^,* BDSC #18576*), Hrb57A (P{EP}HnRNP-K^G13574^,* BDSC #29672*), Hrp59 (P{GSV3}rump^GS6029^,* DGRC Kyoto #200852*), Srrm1 ^B103^* (*SRm160 ^B103^*, (Fan et al. 2014) and *Srrm2340*^Δ*N*^. Since the transposon used for mutagenesis are large (∼10 kb), inserts in the transcribed part were considered to be null alleles, while inserts in promoter regions were considered hypomorphic alleles. If inserted in an intron transposons disrupt splicing, if inserted in the 5’UTR they will prevent translation of the ORF, or if inserted in the ORF lead to a truncated non-functional protein. If inserted in the promotor region, transposon inserts reduce transcription. Whether lethality of mutants mapped to the locus was tested by crossing to chromosomal deficiencies.

The null-allele *Srrm234*^Δ*N*^ (*CG7971*) was generated by *GenetiVision* CRISPR gene targeting services. The 3.2kb deletion at the N-terminus of the gene was generated using sgRNAs AGTCTGCTGGGGACACTGCT and CGCCGCAGGACATATAACAG together with donor template harbouring two homology arms flanking a *loxP 3xP3-GFP loxP* cassette. Left and right homology arms of the donor were amplified using primers CG7971-LAF1 (GTTCCGGTCTCTTAGCCCTGCAGCAGCTTCTGCTTG) and CG7971-LAR1 (TCCAAGGTCTCACAGTTTATATGTCCTGCGGCGCTGC), and CG7971-RAF2 (GTTCCGGTCTCTGTCAGCTGGGAGCCGGCAGTGC) and CG7971-RAR2 (TCCAAGGTCTCAATCGAGTGGAGAACCCATACGTACTTAGATCC), respectively. Successful deletion and integration of the cassette was validated by PCR and Sanger sequencing using primers CG7971-outF1 (CATCGATTGTGTTGCATGAAGTTCAC) and CG7971-outR2 (GGGGAGTATCTGTGAGCAGTTGTATC), and LA-cassette-R (AAGTCGCCATGTTGGATCGACT) and Cassette-RA-F (CCTGGGCATGGATGAGCTGT), respectively. For the analysis of *Dscam* alternative splicing the *3xP3-GFP* marker was removed by Cre mediated recombination using an insert on a third chromosome balancer (*TM6B, P{w*[*+mC*]*=Crew}DH2, Tb*, BDSC #1501) and the resulting chromosome was rebalanced with a zygotically YFP-expressing balancer (*TM6B, P{Dfd-EYFP}3 Sb, Tb*, BDSC #8704) to collect the embryonic lethal homozygous mutants.

For gain of function experiments the following UAS lines, gene switch vector inserts and EP lines were used: *UAS GFP* control line (Gabut et al. 2007), *UAS GFP-X16* (Gabut et al. 2007), *UAS RSF1* (Labourier et al. 1999), *UAS GFP-SC35* (Gabut et al. 2007), *UAS GFP-SF2* (Gabut et al. 2007), *Srrm1 (P{EP}Srrm1^G18603^,* BDSC #26938*), UAS GFP-B52* (Gabut et al. 2007), *Hrp36 (P{GSV6}^GS15926^,* DGRC Kyoto #206416), *Hrp38 (P{GSV6}^GS12795^,* DGRC Kyoto #204283), *Hrp48 ({EPgy2}Hrb27C^EY12571^*, BDSC #20758), *Hrp59 (P{GSV3}^GS6029^,* DGRC Kyoto #200852).

*Hrp36* and *B52* genes lie next to each other. The *Hrp36/B52*^Δ*1*^ double mutant was generated by FRT/FLP mediated recombination using *PBac{RB}e01378* and *PBac{WH}f01884* transposon insertions as described (Zaharieva et al. 2015). The lethal *Hrp36/B52*^Δ*1*^ allele was balanced and validated using primers e01378 Rev (GCCACATTTAGATGATTCAGCATTAT), f01884 Rev (GATTCCAATAGATCCCAACCGTTTCG) and RB 3’ MINUS (TCCAAGCGGCGACTGAGATG).

Lethal lines were rebalanced with balancers expressing YFP zygotically, but not maternally under a *Dfd* promoter (*CyO, P{Dfd-EYFP}2*, BDSC #8578, *TM6B, P{Dfd-EYFP}3 Sb, Tb*, BDSC #8704) to allow for selection of homozygous lethal mutants. Non-GFP expressing 14-18 h embryos were further selected according to the morphology of the auto-fluorescing gut to distinguish them from homozygous balancer carrying animals, which die before they express GFP (Haussmann et al. 2008). For over-expression a third chromosomal *elavGAL4* insert was used (*P{w*[*+mmC*]*=GAL4-elav.L}3*, BDSC #8760).

### RNA extraction, RT-PCR, restriction digestion, denaturing acrylamide gels

Total RNA was extracted using Tri-reagent (SIGMA) and reverse transcription was done with Superscript II (Invitrogen) as described (Koushika et al. 1999) using primer *Dscam* 11RT1 (CGGAGCCTATTCCATTGATAGCCTCGCACAG, 1 pmol/ 20 µl reaction). PCR to amplify *Dscam* exon 9 cluster was done using primers 8F1 (GATCTCTGGAAGTGCAAGTCATGG) and 10R1 ΔST (GGCCTTATCGGTGGGCACGAGGTTCCATCTGGGAGGTA) for 37 cycles with 1 µl of cDNA. Primers were labeled with ^32^P gamma-ATP (6000 Ci/ mmol, 25 mM, Perkin Elmer) with PNK (NEB) to saturation and diluted as appropriate. From a standard PCR reaction with a ^32^P labelled forward primer, 10–20% were sequentially digested with a mix of restriction enzymes (NEB) according to their buffer requirements and temperatures. PCR reaction and restriction digests were phenol/CHCl_3_ extracted, ethanol precipitated in the presence of glycogen (Roche) and analyzed on standard 6% sequencing type denaturing polyacrylamide gels. After exposure to a phosphoimager (BioRad), individual bands were quantified using ImageQuant (BioRad) and inclusion levels for individual variable exons were calculated from the summed up total of all variables. Statistical analysis was done by one-way ANOVA followed by Tukey– Kramer post-hoc analysis using Graphpad prism. Percent inclusion levels of exon 9 variables of embryos were calculated from the total sum of variables. RNA in situs were obtained from flybase as described (Haussmann et al. 2008).

### Sequence analysis

The in silico analysis for SELEX motif occurrence was done by plotting the scores from sliding window using a position weight matrix of the *Drosophila melanogaster* consensus sequence for a given RNA binding protein to the *Dscam* exon 9 variable cluster (p<0.05, Korhonen et al. 2009, Blanchette et al. 2009). The CLIP data was downloaded from GEO as deposited in Blanchette et al. 2009, and the TiMAT window-scores have been used to generate the plot based on the version 4 of the genome assembly. Tiling arrays scores within the *Dscam* variable exon 9 cluster were plotted against all scores to determine whether the four Hrp proteins show significantly increased binding.

Vista alignments were generated as described (Haussmann et al. 2011). The exon 9 docking site was scanned against all gene sequences (as downloaded from FlyBase on the 26^th^ of February 2019) using Blat (Version 35, parameters:-stepSize=1 tileSize=6-minScore=0-minIdentity=0, filtering for hits of at least bit-score of 30 and a coverage over the docking sequence of at least 20 residues) (Kent 2002).

In order to compare the docking site’s propensity to form potential long-range base-pairing with other intronic *Dscam* sequences of the same length, we sampled 100 strictly intronic sequenced of the same or higher sequence complexity as determined by Shannon entropy. We then used the BioPython Bio.pairwise2 local alignment implementation to get pairwise alignments against all (excluding the sampled sequenced and the docking sequence) reverse and reverse-complement sub-sequences of the same length in any *Dscam* intron, using a custom substitution matrix to allow for G-U pairings (scoring scheme: G->U 0.8, A->U: 1, G->C: 1.2, gap opening penalty 0.1, gap extension penalty 0.1) (Cock et al. 2009). We then retained all hits with less than 3 gaps and normalized the alignment scores by each queries’ potential highest score. To assess the best potential secondary structure formed by the docking sequence with any intronic sequence, we run the same sampled sequenced against the concatenated reverse and reverse complement intronic sequences of *Dscam* (masking each query) using RNAduplex from the Vienna package and obtained for each sequence the predicted lowest energy secondary structure prediction (Lorenz et al. 2011).

## Acknowledgments

We thank Bloomington, Kyoto and Harvard stock centres, and J. Tazi and L. Rabinow for fly lines, and P. Grzechnik for comments on the manuscript. We acknowledge funding from the Sukran Sinan Memory Fund to P.U, BBSRC and the European Research Council (ERC-StG-LS2-637591) to M.I.

## Author contributions

M.S. and P.U. designed and directed the project. P.U., M.S., I.H., H.L. performed experiments. A. T-M and M.I. generated and validated the *Srrm234*^Δ*N*^ allele. R.A. performed bioinformatic analysis. P.U. and M.S. wrote the manuscript. All authors read and approved the final manuscript.

## Figure legends for Supplementary Figures

**Supplementary Figure S1.**
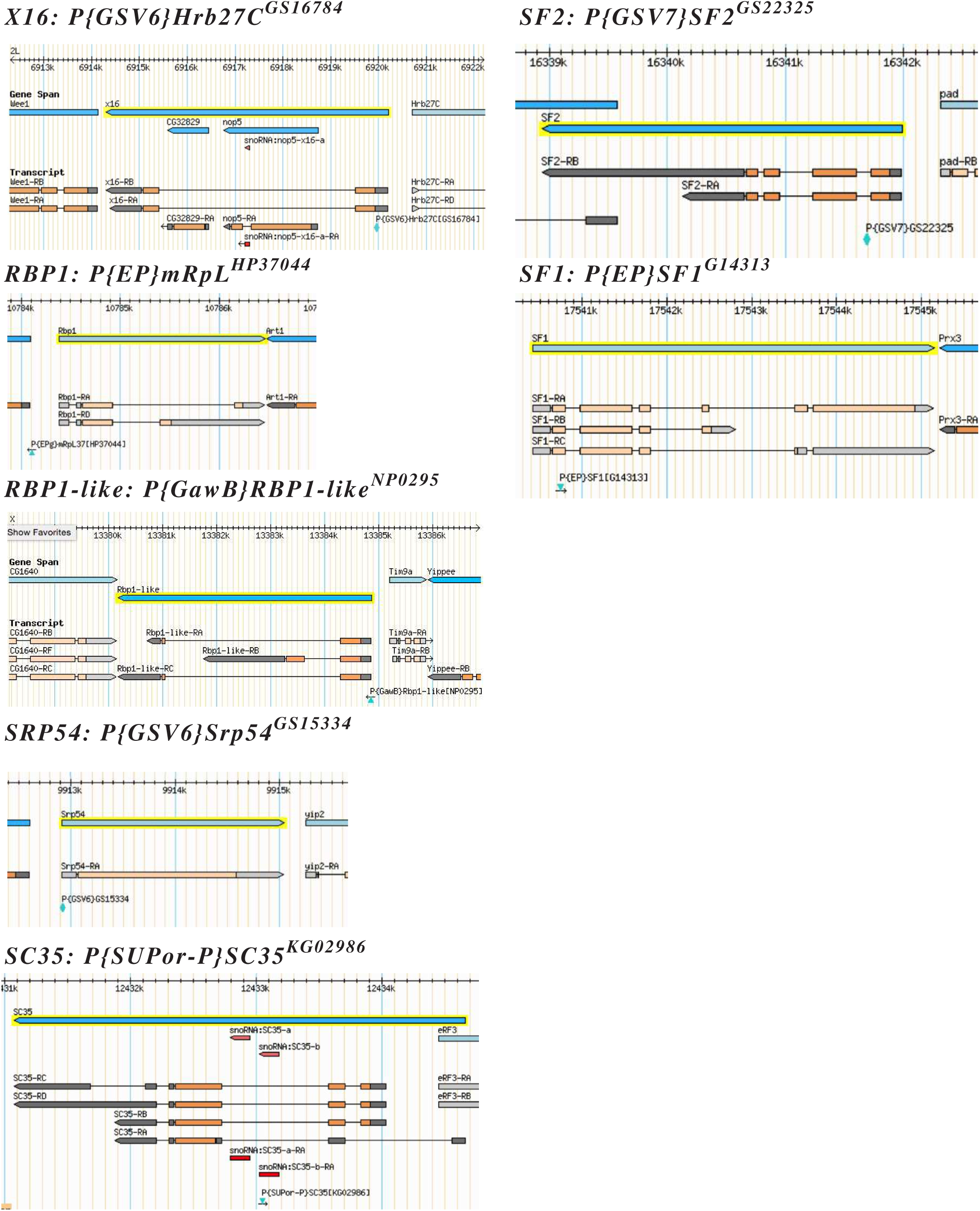
Transposon inserts in SR protein genes. The genomic locus is depicted for the genes where transposon insertion alleles were analysed. The gene span is shown on top as blue box and below transcribed parts are shown as boxes for exons and lines for introns. Transposon insertions are shown as green triangles. Note that transposon have a size of ∼10 kb and disrupt gene expression.

**Supplementary Figure S2.**
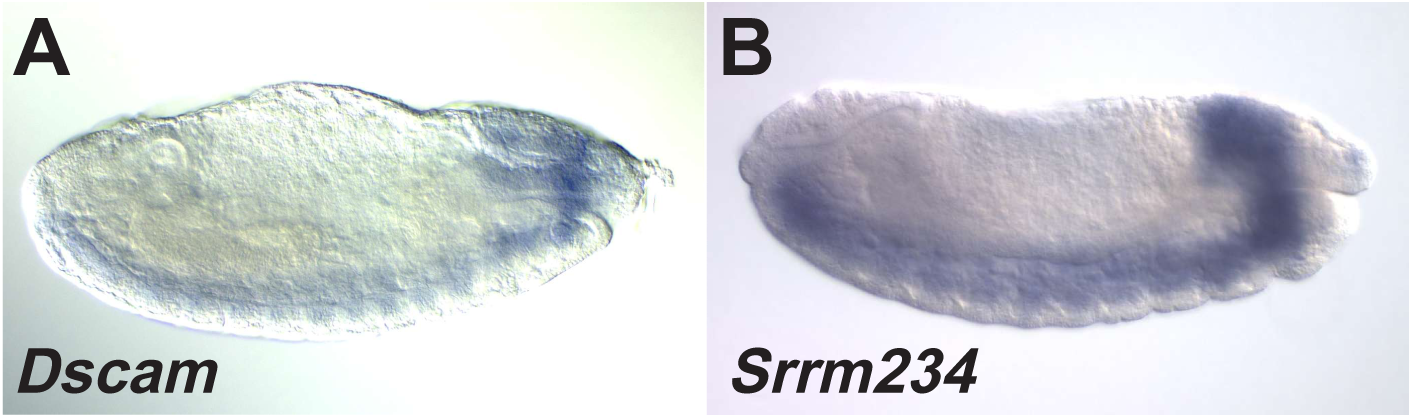
Expression of *Dscam* (A) and *Srrm234* (B) in the nervous system of *Drosophila* late stage embryos as determined by RNA in situ hybridization.

**Supplementary Figure S3.**
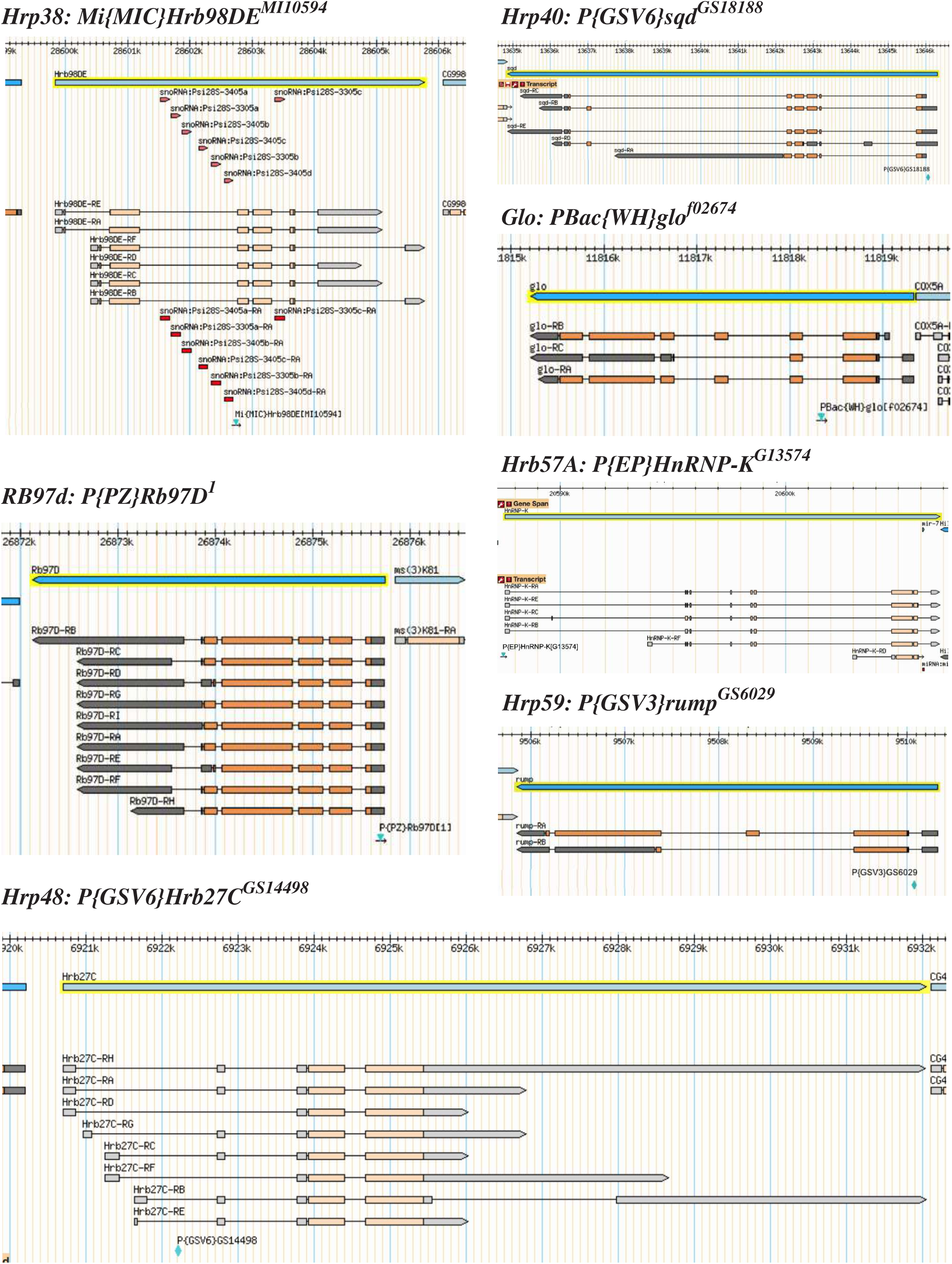
Transposon inserts in hnRNP protein genes. Transposon inserts in SR protein genes. The genomic locus is depicted for the genes where transposon insertion alleles were analysed. The gene span is shown on top as blue box and below transcribed parts are shown as boxes for exons and lines for introns. Transposon insertions are shown as green triangles. Note that transposon have a size of ∼10 kb and disrupt gene expression.

**Supplementary Figure S4.**
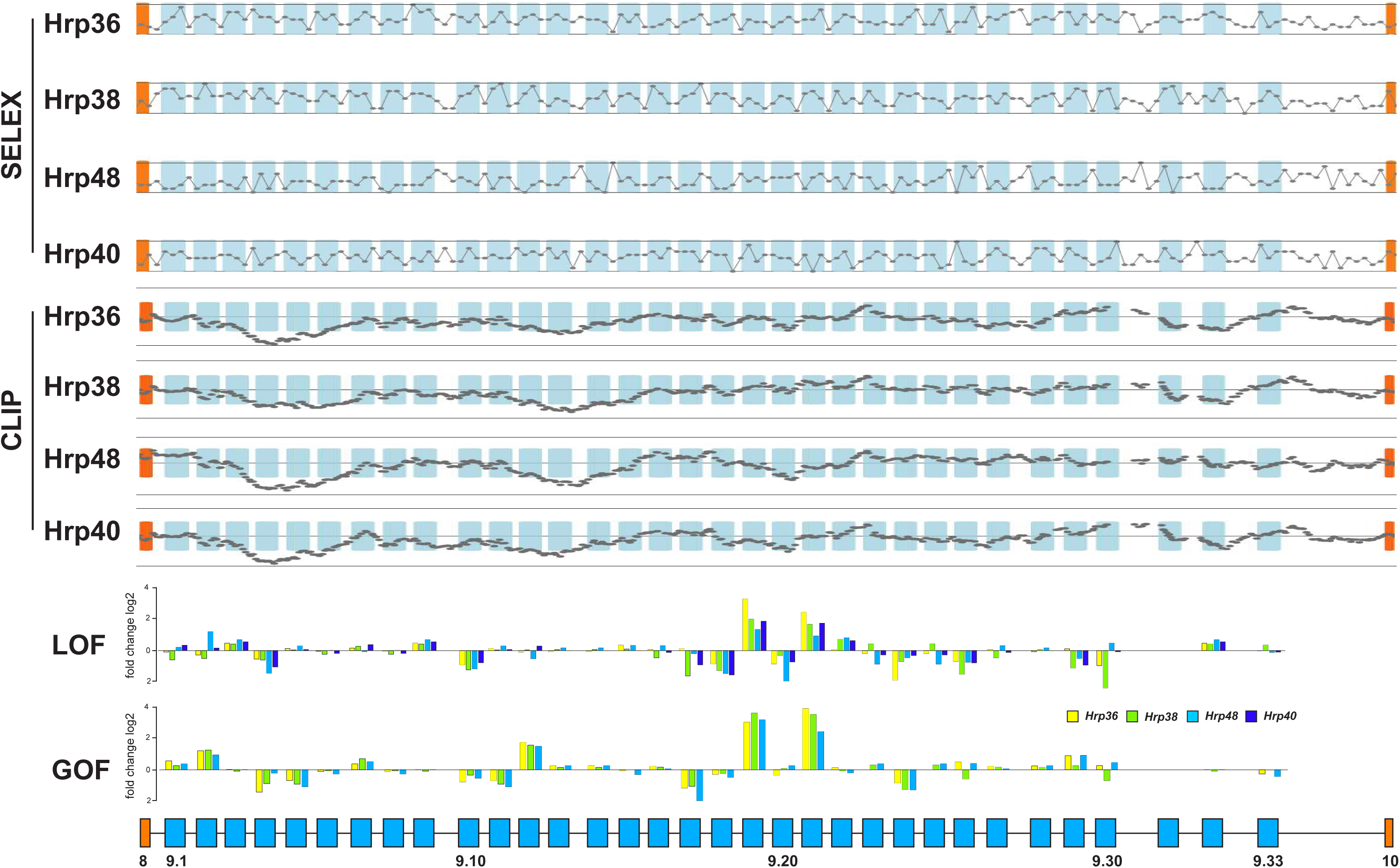
Hrp36, Hrp38, Hrp48 and Hrp40 SELEX motif occurrence is shown for the *Dscam* variable cluster 9 with the line indicating maximal amount of significant hits (p<0.05) of the SELEX motif in a sliding window (12 for Hrp36, 12 for Hrp38, 8 for Hrp48 and 9 for Hrp40). Hrp36, Hrp38, Hrp48 and Hrp40 CLIP binding data from Blanchette et al. (2009) is shown for the *Dscam* variable cluster 9 with top and bottom lines indicating the binding score per tiling oligo probe (±0.54 for Hrp 36 and Hrp38, and ±49 for Hrp48 and Hrp40) compared to averaged binding score for all tiling oligos as determined by TiMAT. Changes in exon inclusion as log2 fold change in LOF and GOF conditions are shown at the bottom.

**Supplementary Figure S5.**
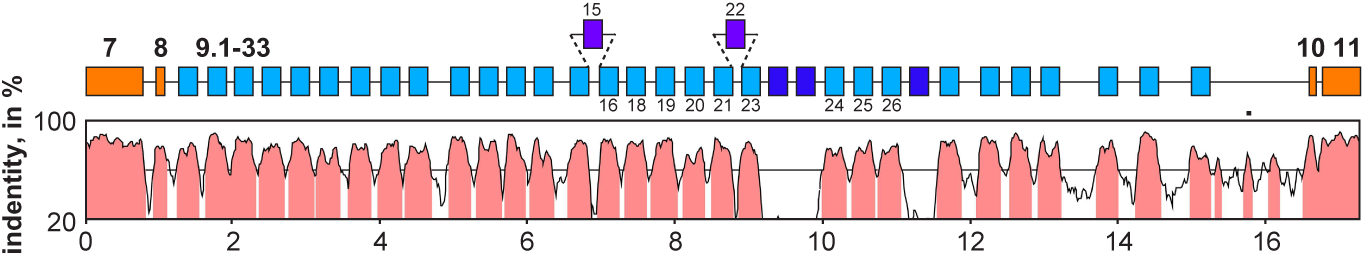
Vista plot alignment of *Dscam* exons 7 to 11 of the *D. melanogaster* sequence compared to the *D. virilis* sequence. Exons are shown as boxes on top and indicated in pink on the alignment. Constant exons are shown in orange and variable exons in light blue. Dark blue exons are absent in *D. virilis*, and exons only present in *D. virilis* are shon on top.

**Supplementary Figure S6.**
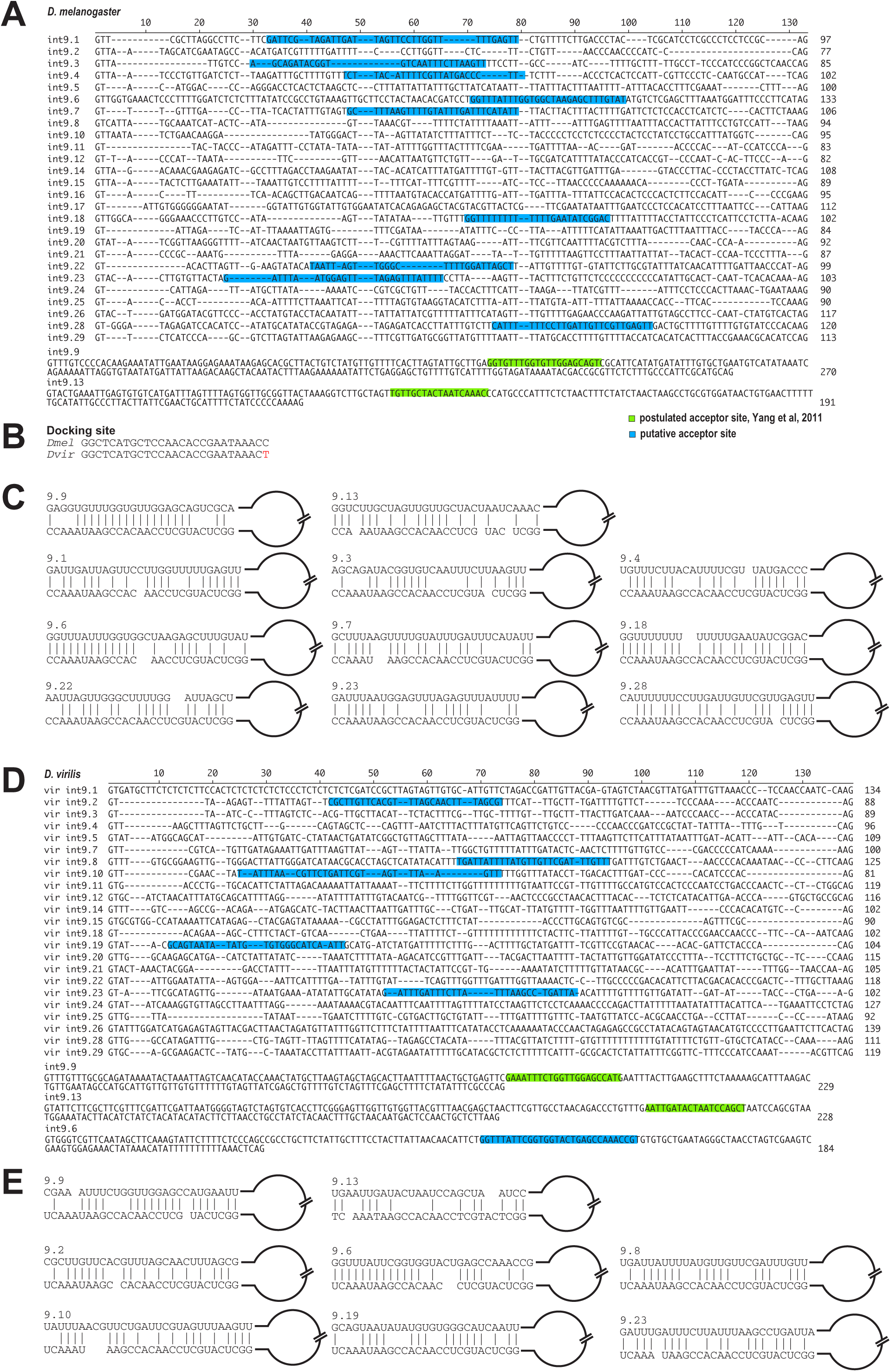
Alignment of the short introns between variable exons from the exon 9 cluster of *D. melanogaster* (A) and *D. virilis* (D), and analysis for sequences complementary to the conserved sequence from intron 9.33 termed docking site (B, Yang et al, 2011). Longer introns with postulated selector sequences are shown at the bottom of the alignment. Sequences with base-pairing capacity to the intron 9.33 sequence termed docking site are shown for *D. melanogaster* (C) and *D. virilis* (E). Note that sequences with base-pairing capacity to the intron 9.33 sequence termed docking site are absent from most introns.

**Supplementary Figure S7.**
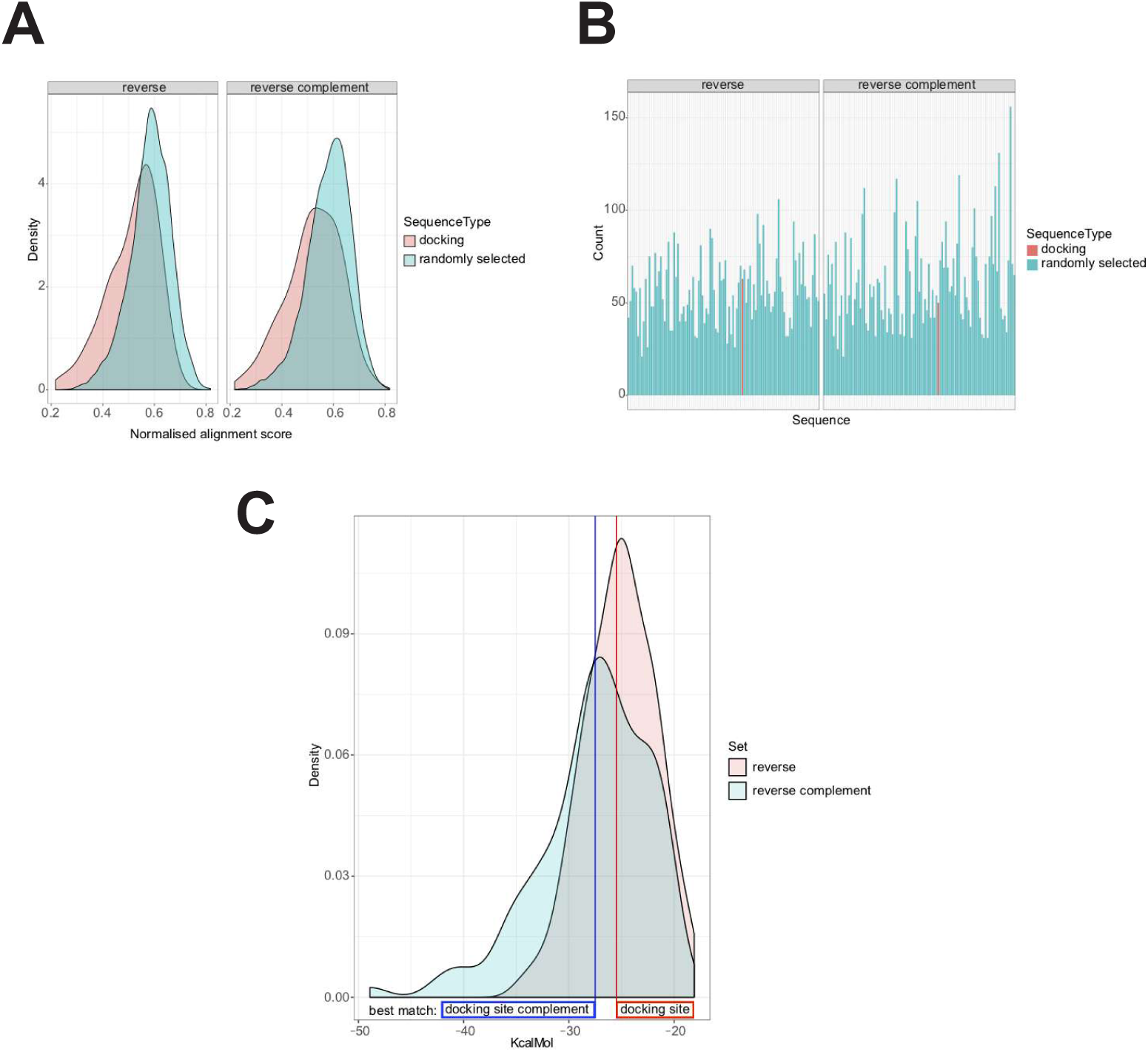
Comparison of long-range base pairing potential of the docking site compared to 100 randomly chosen *Dscam* intronic sequences of the same length. (A) The distribution density of normalised alignment scores (filtered to have at maximum two gaps) against reverse and reverse complement sequences of the same length (docking: alignment scores for the docking sequence, randomly selected: 100 other sequences, see Material and Methods). (B) The same data, but showing the amount of sequences that pass the gap filtering threshold in both sets. (C) Density distribution of computed optimal secondary structures for each sequence. Vertical lines indicate the result for the docking sequence against the reverse and reverse-complement concatenated intronic sequences.

